# The Genotype Diversity within the H5N8 Influenza A Subtype

**DOI:** 10.1101/492470

**Authors:** Andrew Dalby, Lorna Tinworth, Munir Iqbal

## Abstract

The H5N8 influenza subtype has been involved in the global spread of the Guangdong variant of the H5 hemagglutinin. The sequence data from all of the complete genomes from the H5N8 subtype was used to analyse the genotype diversity. Clustering analysis was used to assign lineages to each of the segments where different lineages were present. The results show that there have been multiple reassortment events both within the Guangdong and non-Guangdong H5 hemagglutinin containing variants. The Guangdong H5 variants can be subdivided further into two sub-lineages 2.3.4.4.A and 2.3.4.4.B that have undergone reassortment in both the far east and the United States.

## Introduction

The H5N8 influenza subtype is of considerable interest because of its potential global impact in spreading the Guangdong variant of the H5 hemagglutinin[1].

The highly pathogenic avian influenza virus (HPAIV) H5 hemagglutinin originated in a goose in Guangdong in 1996 [2]. This lineage has subsequently been classified into subgroups/clades by the World Health Organisation [3,4]. This H5-HPAIV nomenclature can be used to examine the diversity of the viral proteins in the H5N8 subtype.

Previously the HPAIV H5 Guangdong lineage had been limited to Asia but the 2014 H5N8 outbreak in Korea was spread through wild bird migration to Western Europe and North America [5–9]. The H5N8 subtype was spread via long range bird migration [10,11]. In North America there was a rapid reassortment that produced the H5N2 subtype which then spread across the US during 2015 [12,13]. Since the winter 2014/2015 European outbreak there have been several subsequent introductions of new isolates that originated in Asia [7,14–16].

Clade 2.3.4.4 of H5N8 subtype can be broken down into at least two distinct groups, clade 2.3.4.4.A and 2.3.4.4.B. These were first identified during the 2014 South Korean outbreak. Previously, these were identified as the Buan (A) and Gochang (B) clades [17,18].

Pairwise distance methods can be used in order to calculate the diversity of the nucleotide or protein sequences and in order to cluster the sequences into groups [19]. Where there is considerable diversity and where the functional properties of the gene are being analysed it makes more sense to use the protein sequence rather than the nucleotide sequence.

Within the sequence alignments there are patterns of conserved and variable amino acids that can be used to define sub-groups for the different proteins. These sub-groups can be identified rapidly by carrying out cluster analysis using the matrix of pairwise distances between sequences [20]. For the H5 hemagglutinin protein these sub-groups can be checked against the existing WHO H5 sub-clade nomenclature [3]. For the other viral proteins there is no currently available classification but it is possible to measure the within group diversity of the clusters to show that the clustering is reliable.

The clusters specify the different lineages and sub-lineages and we can use these to assign a genotype to the particular H5N8 virus.

## Materials and Methods

The complete set of H5N8 protein sequences for all of the viral proteins were downloaded from the Influenza Research Database [21]. These sequences were downloaded as FASTA files with a customized header that included;

Accession

Protein

Subtype

Date

Country

Host

H5 Clade

Strain

Only complete length sequences were used.

Sequences were aligned using Muscle within Mega 7.0.26 [22,23]. The alignment was then visually checked for missing amino acids and for truncated sequences. Any sequences that were not full length were removed, as were any sequences which contained a large number of unassigned amino acids. A summary of the dataset used in the analysis is given in table 1.

**Table One:** Summary of the dataset used for analysis.

| Protein | HA | M1 | M2 | NA | NP | NS1 | NS2 | PA | PB1 | PB2 |
| --- | --- | --- | --- | --- | --- | --- | --- | --- | --- | --- |
| Number of Sequences | 236 | 219 | 205 | 218 | 216 | 199 | 207 | 217 | 205 | 216 |

Pairwise sequence distances were calculated within Mega from the final set of aligned sequences. Distances were calculated using the Poisson correction model within Mega7. Distances are given as the number of amino acid substitutions per site. Excel was used to calculate the mean and maximum pairwise sequence distances for each protein.

Mega produces only the triangular matrix of the pairwise distances. In order to create a complete distance matrix the matrix was imported into R, after removing the sequence names to a separate file in Excel, and the complete matrix generated [24]. From the complete distance matrix a k-means clustering analysis was carried out in R, by firstly finding the number of clusters present in the data. This was determined by calculating the within group sum of the square distances for 2-14 clusters and finding where this falls to a minimum and then levels off. The k-means clustering can then be carried out for the optimum number of clusters and the clustering can used to annotate the pairwise distance matrix. The R-scripts used to carry out the analysis are available as supplementary materials (script files S1-S10). Finally the names of the sequences were added back to the data analysis within Excel.

The genotypes for H5N8 strains were assigned manually based on the clustering. Each genotype corresponds to a unique combination of cluster numbers for the different segments. Where there are segments missing these are provisionally assigned to a distinct genotype dependent on the other cluster numbers being distinctive.

## Results

The maximum, mean, median and variance in the amino acid distances between sequences are given in table 2.

**Table 2:** The summary statistics for the percentage pairwise distances between the sequences for each protein.

| Protein | HA | M1 | M2 | NA | NP | NS1 | NS2 | PA | PB1 | PB2 |
| --- | --- | --- | --- | --- | --- | --- | --- | --- | --- | --- |
| Maximum | 16.40% | 7.00% | 14.40% | 19.00% | 4.30% | 44.60% | 25.20% | 4.90% | 3.40% | 4.60% |
| Median | 2.40% | 1.20% | 1.00% | 1.90% | 0.60% | 1.30% | 2.50% | 1.60% | 1.30% | 1.20% |
| Average | 3.36% | 1.72% | 3.00% | 3.11% | 0.70% | 5.05% | 4.00% | 1.67% | 1.19% | 1.48% |
| Variance | 0.16% | 0.04% | 0.12% | 0.17% | 0.00% | 0.64% | 0.20% | 0.02% | 0.01% | 0.02% |

HA (hemagglutinin), NA (neuraminidase), M1 (matrix-1), M2 (matrix-1), NP (nucleoporin), NS1 (nonstructural-1), NS2 (nonstructural-2), PA (polymerase acidic), PB1 (polymerase basic-1) PB2 (polymerase basic-2).

The number of clusters identified for each protein are given in table 3.

**Table 3:** The number of clusters for each of the proteins.

| Protein | HA | M1 | M2 | NA | NP | NS1 | NS2 | PA | PB1 | PB2 |
| --- | --- | --- | --- | --- | --- | --- | --- | --- | --- | --- |
| Number of Clusters | 3 | 3 | 3 | 6 | 1 | 3 | 7 | 3 | 2 | 2 |

HA (hemagglutinin), NA (neuraminidase), M1 (matrix-1), M2 (matrix-1), NP (nucleoporin), NS1 (nonstructural-1), NS2 (nonstructural-2), PA (polymerase acidic), PB1 (polymerase basic-1) PB2 (polymerase basic-2).

The plots of the within group sum of the squares against the number of clusters are shown for the hemagglutinin (HA), neuraminidase(NA) and nucleoporin (NP) proteins in figures 1-3 respectively.

**Figure 1:**
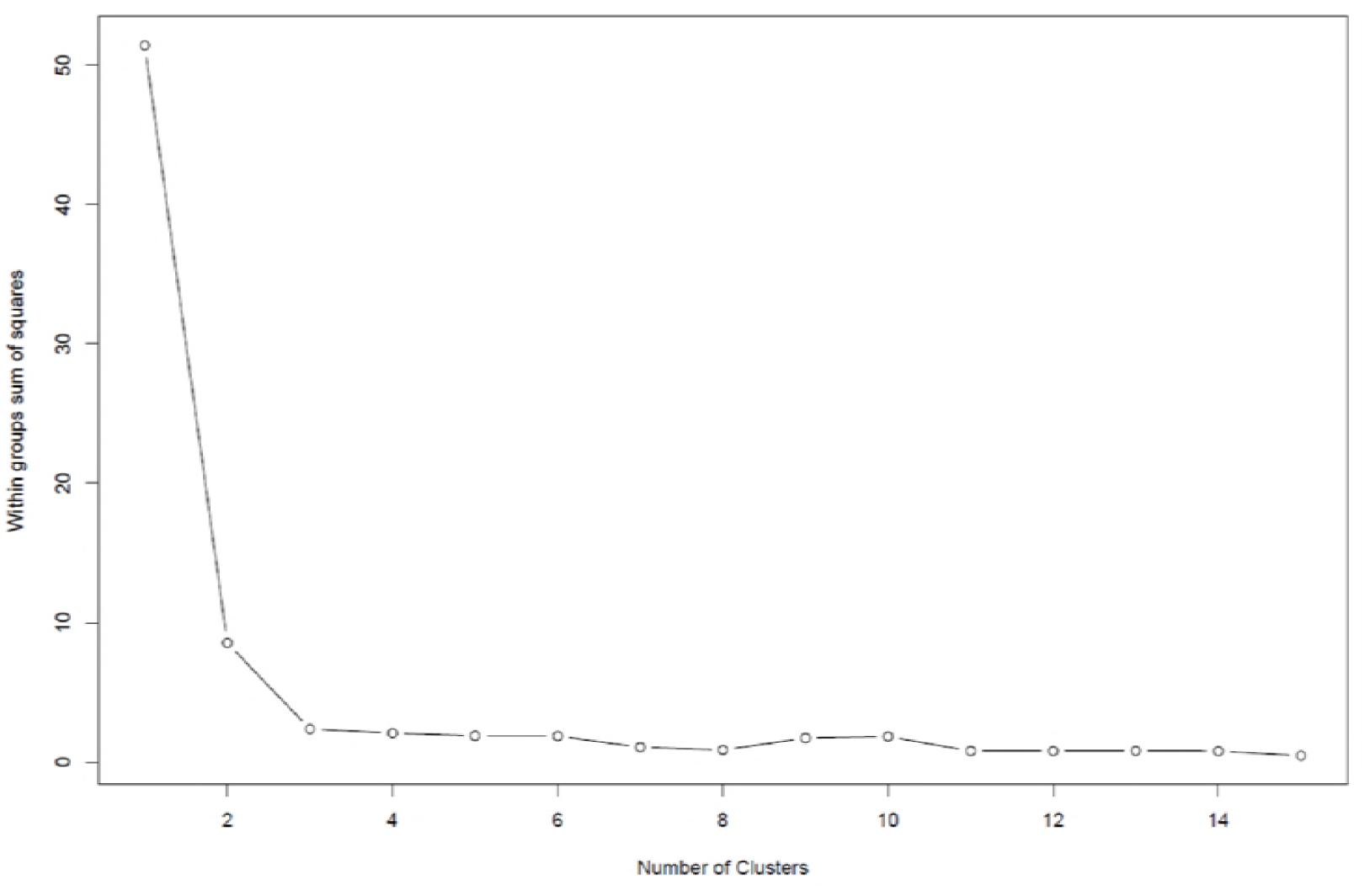
The within group sum of the squares for the hemagglutinin protein.

**Figure 2:**
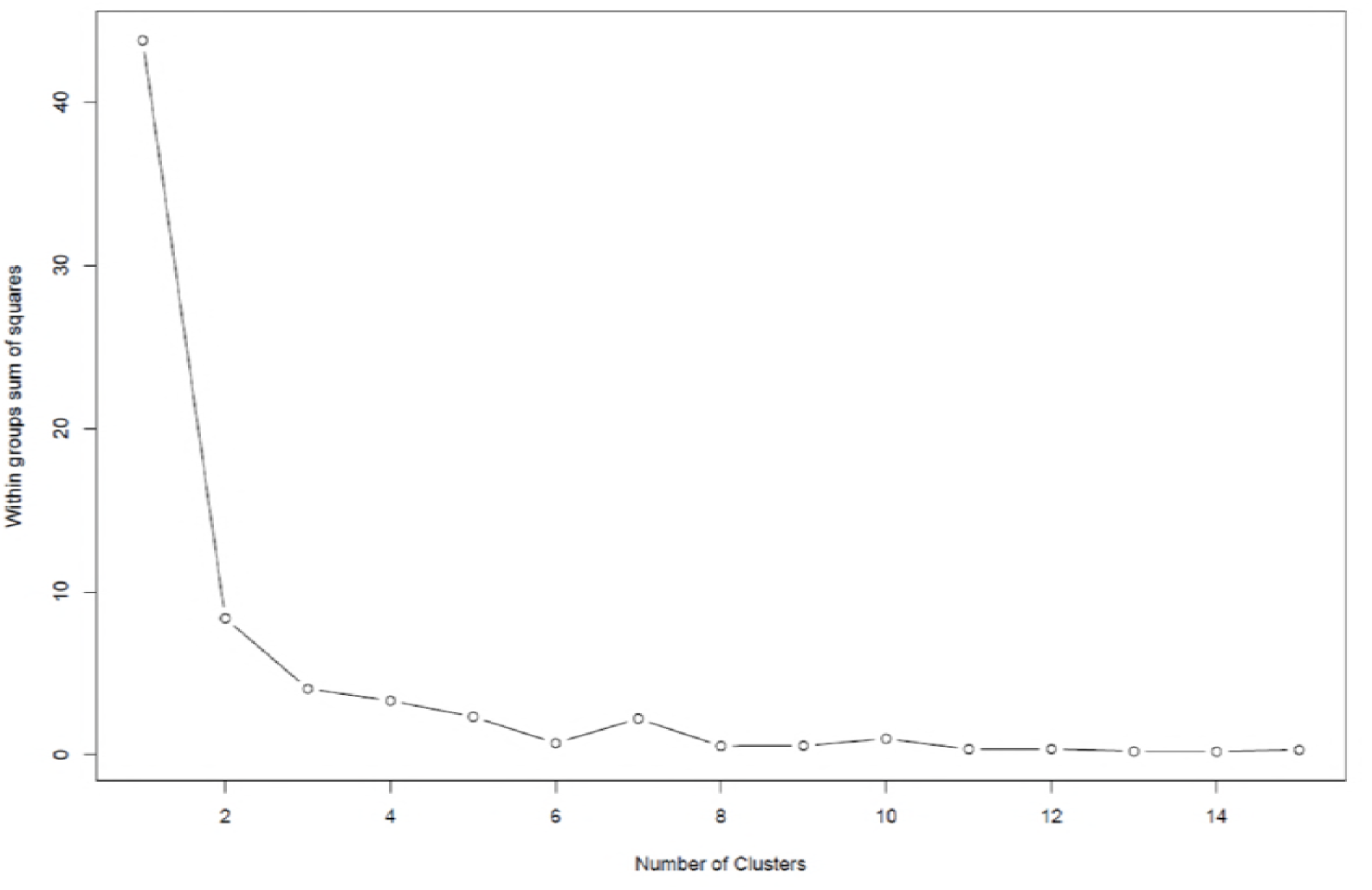
The within group sum of the squares for the neuraminidase protein.

**Figure 3:**
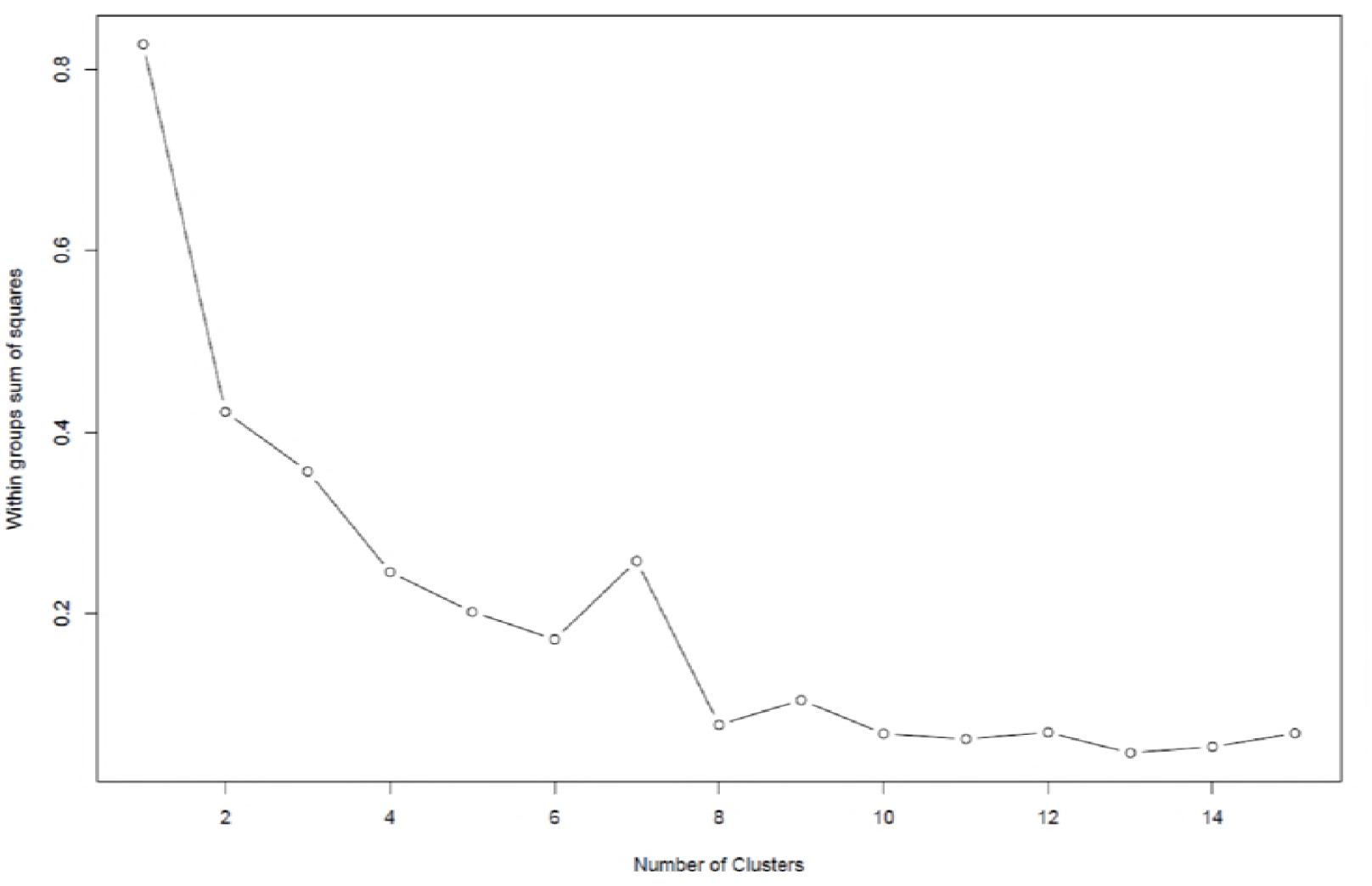
The within group sum of the squares for the nucleoporin protein.

The plots of within group sums of the squares for the other proteins can be found in supplementary materials figures S11-S17

The cluster numbers were assigned to the sequences for each of the different segments. These clusters represent different lineages of the viral protein segments. From cluster assignment the unique genotypes were determined. Each genotype represents a different combination of clusters (lineages) for the different viral protein segments.

The 31 complete or almost complete genotypes are shown in Table 4 along with their representative strains. Representative strains are the first occurrences of that specific combination of viral protein segment lineages.

**Table 4:**
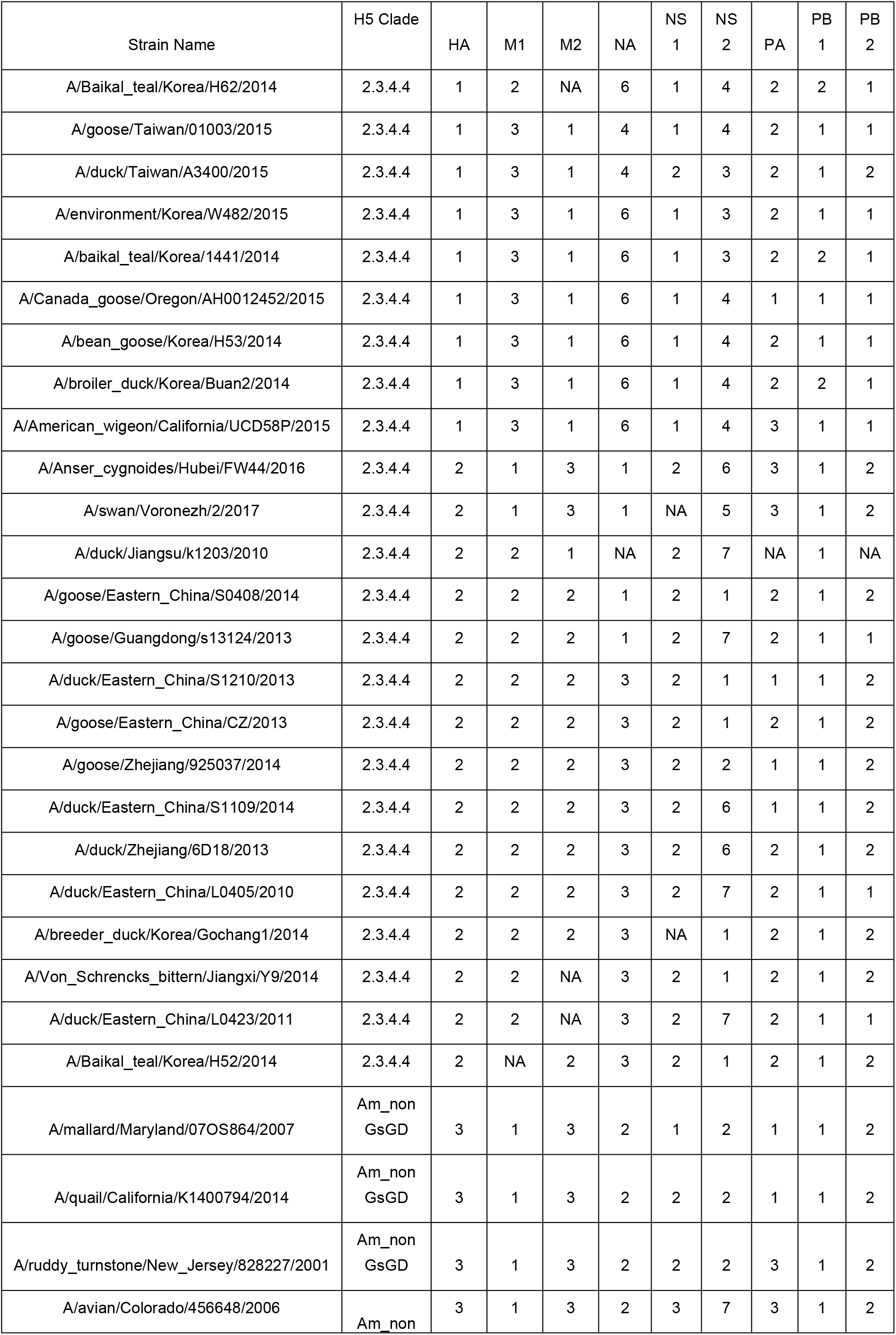
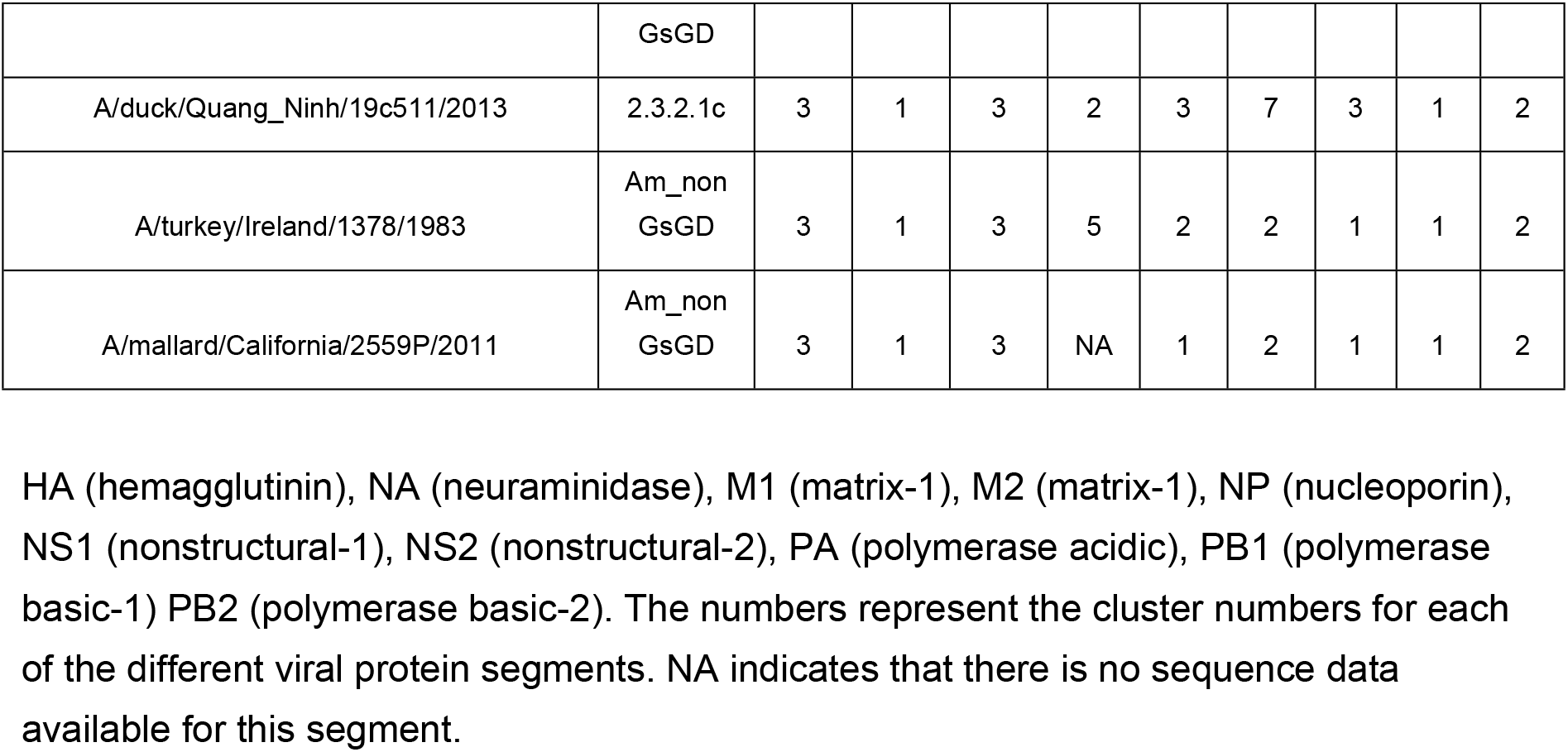
The unique H5N8 genotypes.

## Discussion

From the summary statistics for the pairwise distances between the sequences it is clear that there is very little variation between the matrix 1 (M1), NP, polymerase acidic (PA), polymerase basic 1 (PB1) and polymerase basic 2 (PB2 proteins. These proteins are well conserved as they are not under the same degree of selection as the other proteins. The polymerase genes need to be conserved in order to form a functional polymerase together. The M1 and M2 proteins are on the same viral segment but evolve at different rates. This suggests that it is more appropriate to classify the virus at the protein rather than the viral segment level and to look at evolution on a gene by gene basis.

The biggest differences are in the non-structural proteins (NS), but these differences are very skewed and this indicates one or two clusters of sequences that are distant from the others. The NS1 protein is responsible for inhibiting cellular gene expression as it is an RNA binding protein [25,26]. The role of the NS2 protein is less clear but it is involved in regulating viral replication and it is also known as the nuclear export protein [27–29]. In all of these roles increased diversity of the non-structural proteins may play an important part in viral evolution.

The HA and NA proteins exhibit very similar amounts of variation along with the M2 protein which forms the ion selective proton channel. The glycoproteins HA and NA are well known for their variability through antigenic drift and antigenic shift [30]. The M2 protein is the target for amantadine and rimantadine based anti influenza drugs and this might explain its higher level of variation compared to the M1 protein.

The number of clusters for the K-means clustering of the protein sequences was determined from the graph of the within group sum of the squares (WGSS) against the number of clusters (figures 1-3). The WGSS is initially high when there is only a single cluster as this reflects the total diversity of the sequences, it then falls sharply as the number of clusters increases as the diversity within clusters falls. Once it reaches a minimum then increasing the number of clusters does not reduce the diversity within the clusters and you are then over-fitting the data.

It is important in looking at the graphs of the WGSS to look at the scale of the y-axis. For the HA and NA proteins the WGSS ranges between 0 and 40 (figures 1 and 2). However, for the NP protein the WGSS is between 0 and 0.8 which is over an order and nearly two orders of magnitude less. This very small WGSS in the case of the NP protein reflects the small average pairwise differences between sequences for this segment and shows that there is very little diversity. This indicates that the NP protein contains only a single cluster. The number of clusters for the other proteins is summarized in table 3 and the graphs of WGSS for each protein are given in supplementary figures S11-S17.

Given the number of clusters for each protein there are 40824 possible genotype combinations, but only 31 unique genotypes are observed. This is not surprising because of the temporal and geographical limits on the possible reassortments. Of these genotypes 24 are within the 2.3.4.4 Guangdong H5 clade and the other 7 are either in the American non-Guangdong (Am_nonGsGD, or 2.3.1.2c Guangdong clades.

The clustering of the H5 HA protein identified 3 different clusters. These correspond to the non-2.3.4.4 (cluster 3) sequences and two clusters within the 2.3.4.4 clade. This sub-division of the 2.3.4.4 clade has been reported previously in the studies of the 2014 South Korean outbreak where the HA sequences were divided into the A (Buan) and B Gochang lineages [17,18]. These correspond to cluster 1 (A/Buan) and cluster 2 (B/Gochang) in the current study. There are 9 Buan genotypes and 15 Gochang genotypes.

The presence of a single cluster for the HA sequences containing non-Guangdong H5 sequences as well as Guangdong sequences coupled with the division of the 2.3.4.4. clades into two distinct clusters shows that there is more variability within the Guangdong 2.3.4.4 clade than there is between Guangdong and non-Guangdong sequences. This suggests that the nomenclature for H5 HA needs to be revisited in order to take the non-Guangdong sequences into account.

The first outbreak of H5N8 was in a goose Ireland in 1983 is found in cluster 3 of the HA clusters. This is the only genotype containing the cluster 5 lineage of the N8 neuraminidase, which indicates that reassortment occurred shortly after the origin of the H5N8 subtype. This cluster contains seven different genotypes, most of which are represented by single sequences, or sequences from a single location and year. This agrees with previous work based on the HA and NA phylogenetic trees that showed that in North America H5N8 occurs sporadically and is formed by the reassortment of an H5 and N8 containing subtype, but that it often does not persist [31,32].

This H5 cluster 3 group also contains the cluster 1 and cluster 3 M1 and M2 lineages, which are also present in two recent genotypes from within the Gochang sub-clade in China/Russia. The Quang Ninh genotype is identical to the avian Colorado 2006 genotype except that it contains a 2.3.1.2c H5 and not an Am_nonGsGD hemagglutinin. This evidence suggests that there is extensive long-range movement of avian influenza within the wild bird population.

Almost all of the Buan genotypes contain the 1.3.1.6.1 pattern for the HA, M1, M2, NA and NS1 genes. There was a significant reassortment of the virus from the Buan group during the 2015 Taiwanese out-break where there are different lineages for the N8 neuraminidase and NS1 proteins [33,34]. All of the other genotype variations are in the NS2 and polymerase genes. These show that new lineages of these internal genes were incorporated as H5N8 migrated to North America, but this only formed a single new lineage. The continuing circulation of the H5 2.3.4.4 sub-clade in North America has been disputed [35,36]. Based on the evidence of the current study there has not been a widespread diversification of the H5N8 H5 2.3.4.4 containing genotypes.

There is much more diversity in the Gochang genotypes than in the Buan genotypes. This clade actually originated in Eastern China in 2010 and there were 7 Chinese genotypes before the Gochang genotype evolved during the 2014 South Korean outbreak. There have only been two post 2014 novel Gochang cluster genotypes, one in Russia and the second in China, both of these contain the M protein variants from the original Irish genotypes. The pattern of reassortment for the NS1, NS2 and polymerase proteins is more complex than expected. The NS2 protein is particularly variable and forms 7 different clusters. The polymerase proteins can be classified into fewer clusters but they combine together in different ways more often than the other proteins.

This study has shown that clades 2.3.4.4.A and 2.3.4.4.B containing variants of the H5N8 subtype have been isolated from wild birds and that these have spread to the domestic flocks. A large number of the genotypes identified in the paper originated in wild migratory birds, such as teal, widgeon, swans and geese. None of the genotypes were originally isolated in chickens and only the original outbreak in Ireland in 1984 can be unambiguously assigned to domestic poultry (turkey). Where ducks are the host it is difficult to distinguish between birds that are farmed and those that are wild.

## Supplementary Materials

Script S1: The R-script for the analysis of the HA sequences, Script S2: The R-script for the analysis of the M1 sequences, Script S3: The R-script for the analysis of the M2 sequences, Script S4: The R-script for the analysis of the NA sequences, Script S5: The R-script for the analysis of the NP sequences, Script S6: The R-script for the analysis of the NS1 sequences, Script S7: The R-script for the analysis of the NS2 sequences, Script S8: The R-script for the analysis of the PA sequences, Script S9: The R-script for the analysis of the PB1 sequences, Script S10: The R-script for the analysis of the PB2 sequences.

Figure S11: The within group sum of the squares for the Matrix 1 protein, Figure S12: The within group sum of the squares for the Matrix 2 protein, Figure S13: The within group sum of the squares for the Non-Structural 1 protein, Figure S14: The within group sum of the squares for the Non-Structural 2 protein, Figure S15: The within group sum of the squares for the polymerase A protein, Figures S16: The within group sum of the squares for the polymerase B1 protein, Figure S17: The within group sum of the squares for the polymerase B2 protein.

## Author Contributions

Conceptualization, A.D. and M.I.; methodology, A.D.; software, A.D.; validation, A.D., L.T. and M.I.; resources, A.D.; data curation, A.D.; writing—original draft preparation, A.D.; writing—review and editing, A.D., L.T. and M.I.; visualization, A.D.

## Acknowledgments

None.

## References

1. Verhagen JH, Herfst S, Fouchier RA. How a virus travels the world. Science. 2015;347: 616–617.

2. Xu X, Subbarao K, Cox NJ, Guo Y. Genetic characterization of the pathogenic influenza A/Goose/Guangdong/1/96 (H5N1) virus: similarity of its hemagglutinin gene to those of H5N1 viruses from the 1997 outbreaks in Hong Kong. Virology. 1999;261: 15–19.

3. Who O. Toward a unified nomenclature system for highly pathogenic avian influenza virus (H5N1). Emerging infectious diseases. 2008;14: e1.

4. Smith GJ, Donis RO, Health/Food WHOO for A, Group AO (WHO/OIE/FAO) HEW. Nomenclature updates resulting from the evolution of avian influenza A (H5) virus clades 2.1. 3.2 a, 2.2. 1, and 2.3. 4 during 2013–2014. Influenza and other respiratory viruses. 2015;9: 271–276.

5. Duan L, Bahl J, Smith GJD, Wang J, Vijaykrishna D, Zhang LJ, et al. The development and genetic diversity of H5N1 influenza virus in China, 1996–2006. Virology. 2008;380: 243–254.

6. Vijaykrishna D, Bahl J, Riley S, Duan L, Zhang JX, Chen H, et al. Evolutionary dynamics and emergence of panzootic H5N1 influenza viruses. PLoS pathogens. 2008;4: e1000161.

7. Poen MJ, Bestebroer TM, Vuong O, Scheuer RD, van der Jeugd HP, Kleyheeg E, et al. Local amplification of highly pathogenic avian influenza H5N8 viruses in wild birds in the Netherlands, 2016 to 2017. Eurosurveillance. 2018;23.

8. Ip HS, Torchetti MK, Crespo R, Kohrs P, DeBruyn P, Mansfield KG, et al. Novel Eurasian highly pathogenic avian influenza A H5 viruses in wild birds, Washington, USA, 2014. Emerging infectious diseases. 2015;21: 886.

9. Dalby AR, Iqbal M. The European and Japanese outbreaks of H5N8 derive from a single source population providing evidence for the dispersal along the long distance bird migratory flyways. PeerJ. 2015;3: e934.

10. Lee D-H, Torchetti MK, Winker K, Ip HS, Song C-S, Swayne DE. Intercontinental spread of Asian-origin H5N8 to North America through Beringia by migratory birds. Journal of virology. 2015;89: 6521–6524.

11. Lycett SJ, Bodewes R, Pohlmann A, Banks J, Bányai K, Boni MF, et al. Role for migratory wild birds in the global spread of avian influenza H5N8. 2016;

12. Bui CM, Gardner L, MacIntyre CR. Highly pathogenic avian influenza virus, Midwestern United States. Emerging infectious diseases. 2016;22: 138.

13. Lee D-H, Bahl J, Torchetti MK, Killian ML, Ip HS, DeLiberto TJ, et al. Highly pathogenic avian influenza viruses and generation of novel reassortants, United States, 2014–2015. Emerging infectious diseases. 2016;22: 1283.

14. Beerens N, Heutink R, Bergervoet SA, Harders F, Bossers A, Koch G. Multiple reassorted viruses as cause of highly pathogenic avian influenza A (H5N8) virus epidemic, the Netherlands, 2016. Emerging infectious diseases. 2017;23: 1974.

15. Pohlmann A, Starick E, Grund C, Höper D, Strebelow G, Globig A, et al. Swarm incursions of reassortants of highly pathogenic avian influenza virus strains H5N8 and H5N5, clade 2.3. 4.4 b, Germany, winter 2016/17. Scientific reports. 2018;8: 15.

16. Guinat C, Nicolas G, Vergne T, Bronner A, Durand B, Courcoul A, et al. Spatio-temporal patterns of highly pathogenic avian influenza virus subtype H5N8 spread, France, 2016 to 2017. Eurosurveillance. 2018;23: 1700791.

17. Jeong J, Kang H-M, Lee E-K, Song B-M, Kwon Y-K, Kim H-R, et al. Highly pathogenic avian influenza virus (H5N8) in domestic poultry and its relationship with migratory birds in South Korea during 2014. Veterinary microbiology. 2014;173: 249–257.

18. Lee Y-J, Kang H-M, Lee E-K, Song B-M, Jeong J, Kwon Y-K, et al. Novel reassortant influenza A (H5N8) viruses, South Korea, 2014. Emerging infectious diseases. 2014;20: 1087.

19. Salemi M, Vandamme A-M, Lemey P. The phylogenetic handbook: a practical approach to phylogenetic analysis and hypothesis testing. Cambridge University Press; 2009.

20. Rousseeuw PJ, Kaufman L. Finding groups in data. Seriesin Probability&MathematicalStatistics 1990 34 (1). 1990; 111–112.

21. Squires RB, Noronha J, Hunt V, García-Sastre A, Macken C, Baumgarth N, et al. Influenza research database: an integrated bioinformatics resource for influenza research and surveillance. Influenza and other respiratory viruses. 2012;6: 404–416.

22. Kumar S, Stecher G, Tamura K. MEGA7: molecular evolutionary genetics analysis version 7.0 for bigger datasets. Molecular biology and evolution. 2016;33: 1870–1874.

23. Edgar RC. MUSCLE: a multiple sequence alignment method with reduced time and space complexity. BMC bioinformatics. 2004;5: 113.

24. Statistical Package R. R: A language and environment for statistical computing. Vienna, Austria: R Foundation for Statistical Computing. 2009;

25. Hale BG, Randall RE, Ortín J, Jackson D. The multifunctional NS1 protein of influenza A viruses. Journal of general virology. 2008;89: 2359–2376.

26. Min J-Y, Krug RM. The primary function of RNA binding by the influenza A virus NS1 protein in infected cells: Inhibiting the 2′-5′ oligo (A) synthetase/RNase L pathway. Proceedings of the National Academy of Sciences. 2006;103: 7100–7105.

27. O’Neill RE, Talon J, Palese P. The influenza virus NEP (NS2 protein) mediates the nuclear export of viral ribonucleoproteins. The EMBO journal. 1998;17: 288–296.

28. Neumann G, Hughes MT, Kawaoka Y. Influenza A virus NS2 protein mediates vRNP nuclear export through NES-independent interaction with hCRM1. The EMBO journal. 2000;19: 6751–6758.

29. Robb NC, Smith M, Vreede FT, Fodor E. NS2/NEP protein regulates transcription and replication of the influenza virus RNA genome. Journal of general virology. 2009;90: 1398–1407.

30. Bouvier NM, Palese P. The biology of influenza viruses. Vaccine. 2008;26: D49–D53.

31. Dalby AR. Complete analysis of the H5 hemagglutinin and N8 neuraminidase phylogenetic trees reveals that the H5N8 subtype has been produced by multiple reassortment events. F1000Research. 2016;5: 2463.

32. Dalby A. From sporadic to global: The changing face of H5N8. PeerJ PrePrints. 2015;3: e1489v1.

33. Huang P-Y, Lee C-CD, Yip C-H, Cheung C-L, Yu G, Lam TT-Y, et al. Genetic characterization of highly pathogenic H5 influenza viruses from poultry in Taiwan, 2015. Infection, Genetics and Evolution. 2016;38: 96–100.

34. Lee M-S, Chen L-H, Chen Y-P, Liu Y-P, Li W-C, Lin Y-L, et al. Highly pathogenic avian influenza viruses H5N2, H5N3, and H5N8 in Taiwan in 2015. Veterinary microbiology. 2016;187: 50–57.

35. Krauss S, Stallknecht DE, Slemons RD, Bowman AS, Poulson RL, Nolting JM, et al. The enigma of the apparent disappearance of Eurasian highly pathogenic H5 clade 2.3. 4.4 influenza A viruses in North American waterfowl. Proceedings of the National Academy of Sciences. 2016;113: 9033–9038.

36. Ramey AM, Spackman E, Kim-Torchetti M, DeLiberto TJ. Weak support for disappearance and restricted emergence/persistence of highly pathogenic influenza A in North American waterfowl. 2016;

